# Identification of a novel basal body-localized epsilon-tubulin in *Leishmania*

**DOI:** 10.1101/2024.09.27.615455

**Authors:** Arunava Seth, Anubhab Das, Rupak Datta

**Author notes:** Corresponding authors: Rupak Datta Arunava Seth. Worked at the Department of Biological Sciences, IISER Kolkata as a summer intern; present address: Department of Ophthalmology, Visual and Anatomical Sciences, Wayne State University School of Medicine, Detroit, MI, USA.

## Abstract

Epsilon-tubulins are unconventional isoforms of the tubulin family, found in only a few organisms so far. We identified a novel epsilon-tubulin in *Leishmania major* (Lme-tubulin) that exhibits significant sequence similarity and conservation of functional domains with its known counterparts. Lme-tubulin was found to be constitutively expressed in both the extracellular promastigote form of the parasite and the amastigotes residing within infected macrophages. For localization studies, we generated a *Leishmania* strain expressing mNeonGreen-tagged Lme-tubulin using CRISPR-Cas9-mediated endogenous protein tagging method. Imaging studies with this strain revealed Lme-tubulin to be localized near the kinetoplast and at the flagellar base, indicating a basal body localization. That Lme-tubulin is indeed localized at the basal body and not a part of the microtubular network was confirmed when its localization was found to remain unaltered upon treatment with nocodazole, a microtubule disruptor. This is the first experimental finding of an e-tubulin not only in the genus *Leishmania* but in the entire *Trypanosomatidae* family and is likely to incite further research to uncover the physiological role of this intriguing tubulin in this group of protozoan parasites.

**Summary Statement:** We report for the first time an epsilon-tubulin in *Leishmania* parasite. This nocodazole-insensitive unconventional tubulin is expressed constitutively and found to be localized in the basal body.

## Introduction

*Leishmania* species are causal agents for leishmaniasis, a group of diseases with diverse clinical manifestations. Depending on the species involved, the symptoms of leishmaniasis can range from fatal infections of the spleen and liver to self-resolving skin lesion (Burza et al., 2018). Despite this diversity of symptoms, all *Leishmania* species share a similar digenetic life cycle involving a sandfly vector, typically from the genus *Phlebotomus* or *Lutzomyia*, and a mammalian host such as humans. During this lifecycle, the parasite switches between two distinct morphological forms. In the insect stage, it appears as a long, slender promastigote with a motile flagellum. After infecting the host macrophages, the promastigotes transform into small, rounded amastigotes with short, non-motile flagellum. This promastigote-to-amastigote transition is crucial for the survival of intracellular *Leishmania*. The drastic morphological changes involved in this process requires substantial cytoskeletal restructuring (Gluenz et al., 2010; Sunter and Gull, 2017).

Tubulins, along with actin, are pivotal in regulating cytoskeleton dynamics. They play crucial roles in various cellular functions such as vesicular trafficking, cell shape maintenance and cell cycle progression in most eukaryotic cells, including lower eukaryotes like *Leishmania* (Dandugudumula et al., 2022; Fourriere et al., 2020; Gull, 1999; Janke and Chloë Bulinski, 2011; Pollard and Cooper, 2009). Microtubule organizing centers (MTOCs) are the primary nucleation sites for tubulin polymerization. The MTOCs exist in different forms, such as basal bodies associated with cilia or flagella, and centrosomes linked to mitotic spindles (Wu and Akhmanova, 2017). In *Leishmania* as well as in other members of the *Trypanosomatidae* family, the basal body is closely associated with the kinetoplast near the flagellar pocket, a site through which processes like endocytosis or exocytosis occur (Field and Carrington, 2009; Overath et al., 1997; Sengupta et al., 1999; Stierhof et al., 1994). It also plays critical role in formation and anchoring of the flagellum thereby acting as controlling center for parasite’s motility, attachment and infectivity (Halliday et al., 2020; Sunter et al., 2019).

The *Leishmania* basal body is reported to form axoneme structure of classical 9+2 arrangement of microtubules in the promastigote form and a 9+0 arrangement in the amastigotes (Alexander, 1978; Rudzinska et al., 1964; Wheeler et al., 2015). Limited investigations on the *Leishmania* basal body composition have so far identified only a few key proteins, such as centrin, centrin4, protein of centriole, and SAS6 (Selvapandiyan et al., 2001; van Breugel et al., 2014; Vats et al., 2023). Apart from these, γ-tubulin is also known to localize in its basal body (Libusová et al., 2004). In contrast, alpha and beta-tubulin of *Leishmania* act as monomers for protofilaments forming microtubules but are not part of the basal body (Fong and Lee, 1988; Wirth and Slater, 1983). However, the presence of other unconventional tubulin isoforms, such as epsilon, delta, and zeta tubulins, remains to be experimentally validated in *Leishmania* as well as in other parasites of the *Trypanosomatidae* family.

The epsilon, delta, and zeta-tubulins, collectively grouped as evolutionary co-conserved ZED module, are predicted to be present in most eukaryotes, ranging from unicellular protists to higher-order mammals (Turk et al., 2015). Interestingly, genome analysis across various organisms revealed a unique pattern of conservation of the ZED module tubulins. It was observed that organisms lacking epsilon-tubulin were also missing delta and zeta-tubulins, while those possessing epsilon-tubulin consistently had delta and/or zeta-tubulin as well. This observation suggests that epsilon-tubulin might be playing a central role in coordinating the function of ZED module, acting in partnership with delta and/or zeta-tubulin (Turk et al., 2015). Epsilon-tubulin was first predicted in *Saccharomyces cerevisiae* by RG Burns through extensive bioinformatic analysis (Burns, 1995). Subsequently, it was discovered in many other organisms, including humans (Chang and Stearns, 2000; Dupuis-Williams et al., 2002; Dutcher et al., 2002; Ross et al., 2013). In human cells, the epsilon-tubulin was found to localize in the centrosome but at a distinct site from γ-tubulin (Chang and Stearns, 2000). Recently, it has been shown that knocking out epsilon-tubulin in human hTERT RPE-1 cells results in the formation of abnormal centrioles lacking triplet microtubules (Wang et al., 2017). Epsilon-tubulin from *Xenopus laevis*, which is similar to its human counterpart in terms of sequence and localization, has been reported to be important for centriole duplication (Chang et al., 2003). In mice, epsilon-tubulin has been shown to be crucial for forming complex microtubule arrays, and its absence resulted in drastic loss of germ cells leading to male sterility (Stathatos et al., 2024). The basal body-localized epsilon-tubulin of *Paramecium* was found to play essential role in the assembly and anchorage of the centriolar microtubules (Dupuis-Williams et al., 2002). In *Tetrahymena thermophila*, deletion or mutation of the epsilon-tubulin gene caused flagellar defects, with some mutations proving to be lethal (Ross et al., 2013). Similarly, in *Chlamydomonas*, deletion of epsilon-tubulin resulted in failure in basal body assembly, preventing proper flagella formation (Dutcher et al., 2002). Despite these studies proving the essentiality of epsilon-tubulin in various organisms across different branches of life, its presence in the *Trypanosomatidae* group of protozoan parasites remains ambiguous. Although Vaughan *et. al.* have bioinformatically predicted the presence of epsilon-tubulin in *Trypanosoma brucei*, its experimental validation is still awaited (Vaughan et al., 2000). Recently, by employing a CRISPR-Cas9 based high-throughput endogenous protein tagging technique, Billington *et. al.* were successful in fluorescent-labeling of 89% of the *T. brucei* proteome and in mapping their subcellular localization. However, for the epsilon-tubulin, no signal above the background fluorescence was detected in their studies (Billington et al., 2023). A similar high-throughput study is underway to endogenously label the *Leishmania* proteome (Aellig et al., 2024). But till date, there is no report on epsilon-tubulin in any of the *Leishmania* species.

We report here identification of an epsilon-tubulin in *Leishmania major* (*L. major*), which exhibited constitutive expression in both the extracellular promastigote and intracellular amastigote forms of the parasite. Using CRISPR-Cas9 mediated endogenous protein tagging, coupled with imaging studies, we demonstrated that this nocodazole-insensitive unconventional tubulin is localized at the basal body. This is the first experimentally validated epsilon tubulin in the entire *Trypanosomatidae* family. This discovery is likely to encourage further structure-function studies aimed at understanding the role of epsilon tubulin in parasite physiology.

## Results and Discussion

### Identification of *L. major* epsilon-tubulin

Genome sequence of *L. major* predicted the presence of a putative epsilon-tubulin on chromosome 21, annotated as LmjF.21.1010 (Ivens, 2005). The predicted protein sequence of *L. major* epsilon-tubulin (Lme-tubulin) was aligned with bona fide, well-characterized epsilon-tubulins from *Tetrahymena thermophila* (Ross et al., 2013), *Chlamydomonas reinhardtii* (Dutcher et al., 2002), *Homo sapiens* (Chang and Stearns, 2000) and *Paramecium tetraurelia* (Dupuis-Williams et al., 2002) (Fig 1). Lme-tubulin was found to share ∼ 40% sequence identities and ∼ 60% sequence similarities with its counterparts in these species (Fig 1, Table S1). InterPro analysis confirmed the presence of signature tubulin domains, the GTPase domain and C-terminal domain, in Lme-tubulin (Fig 1; yellow and green shaded regions) (Nogales et al., 1998a; Nogales et al., 1998b; Paysan-Lafosse et al., 2023). Importantly, 23 out of 27 GTP-binding residues of *Tetrahymena* epsilon-tubulin are either conserved or replaced with similar residues in Lme-tubulin (Fig 1; black triangles). Also, nine out of ten amino acids, mutations of which in *Tetrahymena* epsilon-tubulin led to lethal or slow-growing phenotypes, were found to be conserved in Lme-tubulin (Fig 1; black boxes) (Ross et al., 2013). Studies with *Paramecium* and human epsilon-tubulins revealed that they have unique amino acid insertions, which are absent in canonical tubulins (Dupuis-Williams et al., 2002; Inclán and Nogales, 2001). These insertions were hypothesized to be required for longitudinal contact with other tubulins (Inclán and Nogales, 2001). We probed for the presence of such insertions in Lme-tubulin by aligning its sequence with a, b and g-tubulins of *L. major* (Fig S1A). Most of the regions aligned except two prominent stretches of amino acids; Insertion I (R211 - T236) and Insertion II (H325 - T332) (Fig S1A). Comparison of the AlphaFold structure of Lme-tubulin with that of the structures of α, β and γ-tubulins of *L. major* showed mostly aligned regions, except for the insertions I and II (Fig S1B). The sequence and structural alignment results thus confirmed the presence of these characteristic insertions in Lmε-tubulin. Interestingly, we observed that the C-terminal of the Lme-tubulin also did not align properly with the α, β or γ-tubulins (Fig S1B). The C-terminal residues of canonical tubulins are reported to be crucial for post-translational modifications (Janke and Chloë Bulinski, 2011). So, a lack of proper alignment and absence of secondary structure of Lmε-tubulin C-terminal region suggests that the post-translational modifications are either absent or different from the classical tubulins. A similar feature was earlier observed in the C-terminal domain of *Tetrahymena* epsilon-tubulin (Ross et al., 2013). Isoforms of Lme-tubulin were also predicted in the genomes of other *Leishmania* species, each displaying very high sequence identity (>97% for *L. donovani*, *L. infantum*, *L. mexicana* and >90% for *L. braziliensis)* (Table S1). Lme-tubulin was found to exhibit >65% sequence identity and >80% sequence similarity with putative epsilon tubulins of other members of the *Trypanosomatidae* family, *T. brucei* and *T. cruzi* (Table S1). Collectively, these findings from our bioinformatic analyses strongly affirm the legitimacy of Lme-tubulin as an authentic epsilon tubulin.

**Fig 1.**
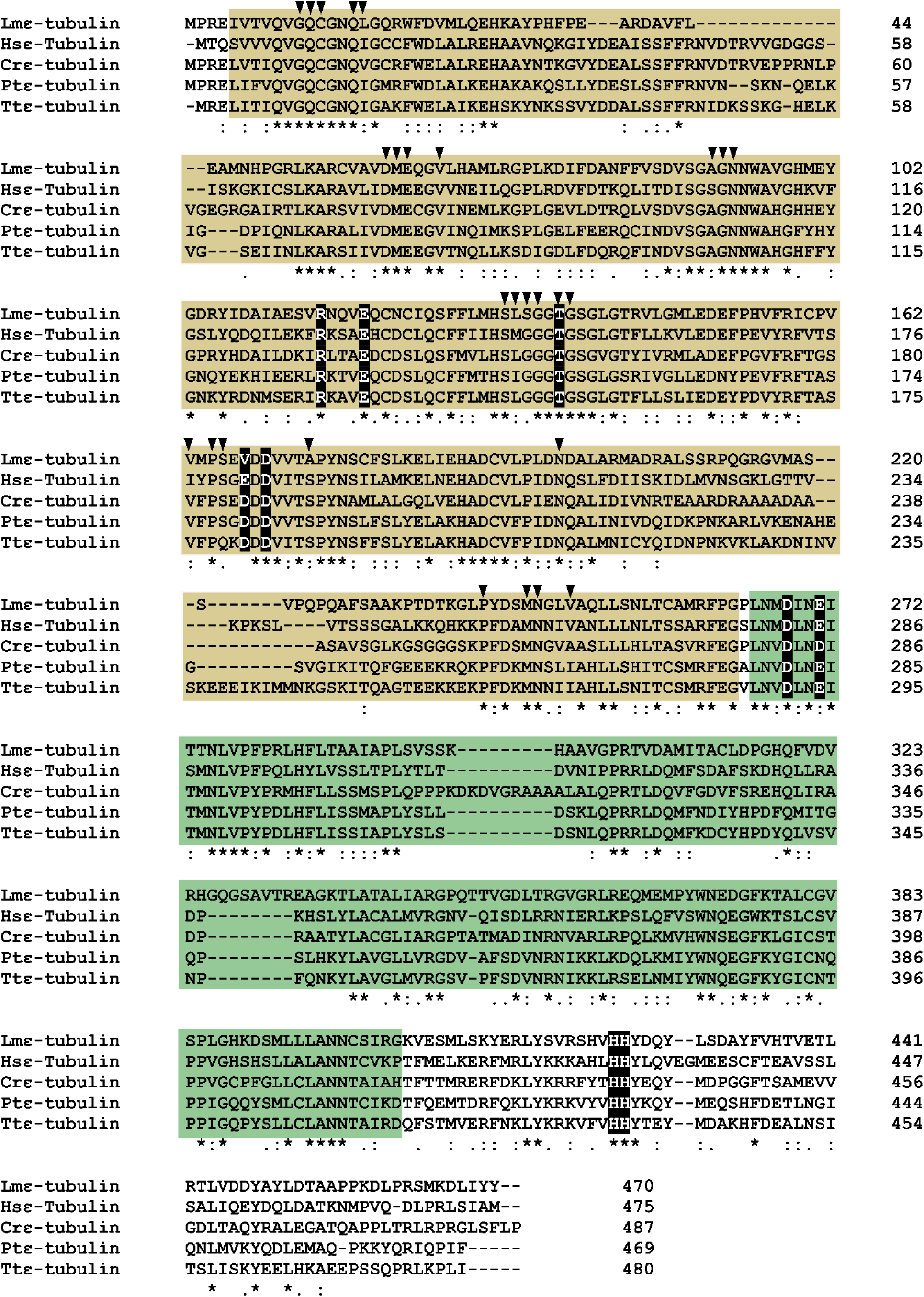
Sequence homology of Lme-tubulin with its counterparts in other organisms. Sequence alignment of Lme-tubulin with epsilon-tubulins from *Homo sapiens* (Hs-e-tubulin) *Chlamydomonas reinhardtii* (Cr-e-tubulin) *Paramecium tetraurelia* (Pt-e-tubulin) *Tetrahymena thermophila* (Tt-e-tubulin). Yellow-shaded region indicates the GTPase domain and green-shaded region indicates the C-terminal domain as predicted by InterPro analysis. Residues marked by black boxes indicate lethal or slow-growing mutations in *Tetrahymena thermophila.* Black triangles indicate GTP-binding residues of tubulins as predicted by InterPro analysis.

### Promastigote and amastigote forms of *L. major* both expresses epsilon-tubulin

Analysis of the genetic locus revealed that the *Lme-tubulin* is flanked upstream by a hypothetical gene (LmjF.21.1000) and downstream by the putative dynein arm light chain encoding gene (LmjF.21.1020) (Fig 2A). To experimentally verify the presence of *Lme-tubulin* in the genome, we used *L. major* genomic DNA as the template for PCR amplification of the corresponding segment using *Lme-tubulin-*specific primers (P1 & P2). A 1413 bp product was obtained, which matched with the expected size of *Lme-tubulin* coding region (Fig 2B). After confirming the presence of *Lme-tubulin* in the genome, our next goal was to check its expression. For this, cDNAs, prepared from *L. major* promastigote mRNAs, were subjected to PCR amplification with primers, P1 & P2. This RT-PCR yielded 1413 bp product, confirming the expression of Lme-tubulin mRNA in the promastigote stage (Fig 2C). Since many genes of *Leishmania* exhibit stage-specific expression pattern, we wanted to verify whether Lme-tubulin is expressed in the intracellular amastigote form also (Brotherton et al., 2010). For this, as described schematically in Fig 2D, we infected J774A.1 macrophage cells with *L. major* promastigotes and isolated total mRNAs from the infected macrophages at ∼ 60 hours post-infection (to allow promastigote-to-amastigote transformation). This pool of mRNA was a mixture derived from both host macrophage cells and the intracellular amastigotes. When RT-PCR was performed with primers P1 & P2 using this mRNA pool from the *Leishmania* infected macrophages, a 1413 bp product got amplified confirming the presence of Lme-tubulin transcripts in the amastigotes. That this product was not a non-specific amplification from macrophage mRNA was confirmed when we found that mRNA isolated from uninfected macrophages did not yield any product upon RT-PCR (Fig 2E).

**Fig 2.**
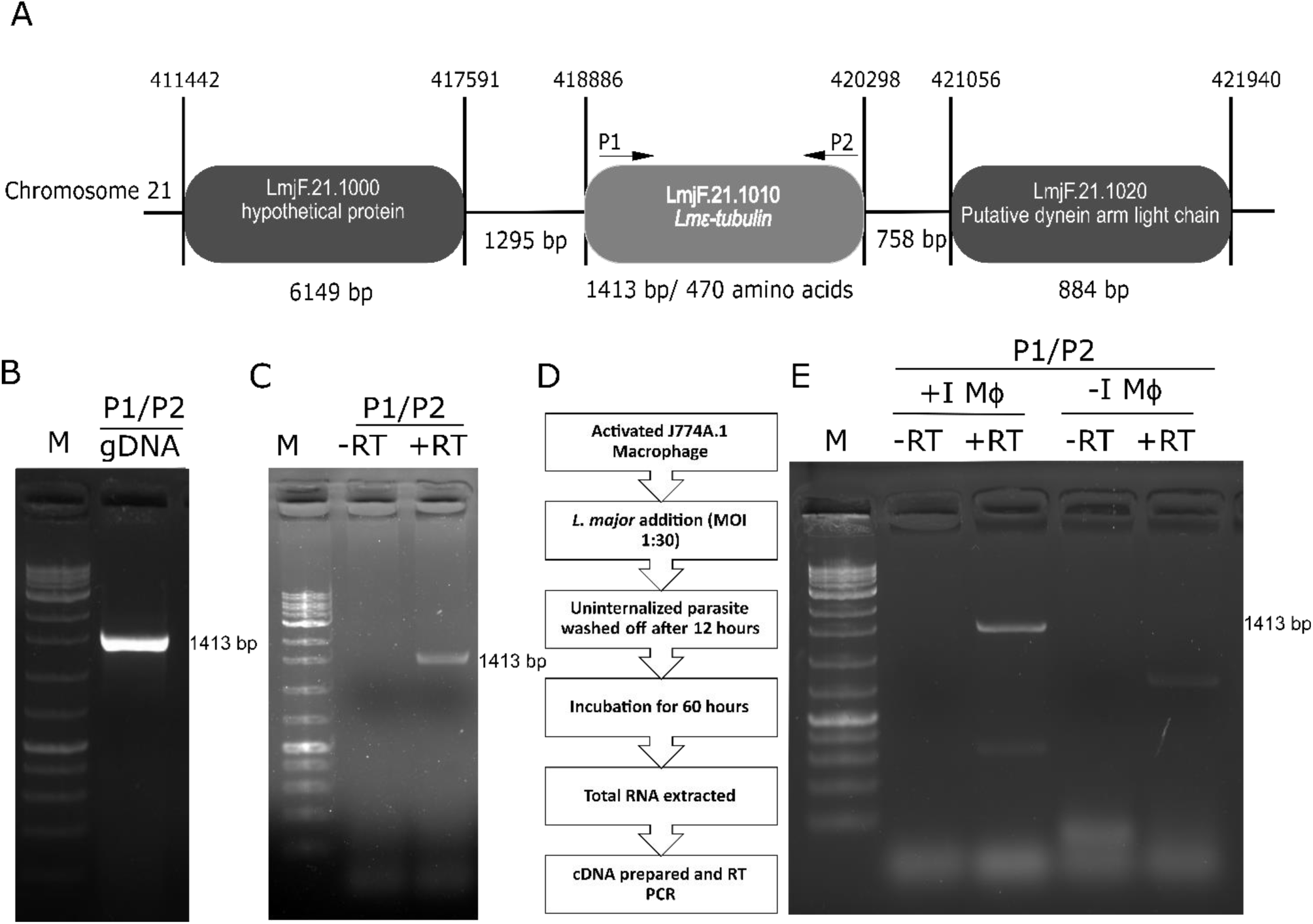
Genomic PCR and RT-PCR analysis of Lme-tubulin. [A] Schematic diagram of the chromosome 21 locus containing *Lme-tubulin* and the adjacent genes. Binding sites of the Lme-tubulin-specific primers (P1/P2) are shown. [B] Agarose gel image showing PCR amplification of 1413 bp *Lme-tubulin* genomic locus with primer sets P1/P2. [C] Expression of Lme-tubulin mRNA in *L. major* promastigotes by RT-PCR using P1/P2. The lanes marked as ‘+RT’ and ‘−RT’ represent the reactions with and without RT enzyme, respectively. [D] Workflow for checking of expression of Lme-tubulin mRNA in *L. major* amastigotes. [E] Expression of Lme-tubulin mRNA in *L. major* amastigotes by RT-PCR using P1/P2. The lanes marked as ‘+RT’ and ‘−RT’ represent the reactions with and without RT enzyme, respectively. +IMΦ and -IMΦ indicates RNA extracted from infected and uninfected macrophages, respectively.

### CRISPR-Cas9 mediated tagging of endogenous epsilon-tubulin with mNeonGreen

Determining the localization of a protein is a prerequisite to understand its function. To examine the localization of Lme-tubulin, we planned to tag the endogenous protein with the fluorophore mNeonGreen using CRISPR-Cas9 gene editing technology (Beneke et al., 2017). For this, we first generated an *L. major* strain expressing *Streptococcus pyogenes* Cas9 (SpCas9) and T7 Polymerase (T7Pol). The strain was verified using genomic DNA PCR, using primers specific for SpCas9 and T7pol. This produced 4.1 kb and 2.7 kb PCR products, respectively, confirming successful integration of SpCas9 and T7pol in the *L. major* genome (Fig S2). This strain, referred to as LmCas9:T7Pol, was used to generate the N-terminal mNeonGreen (mNG)-tagged-Lme-tubulin expressing *L. major* (Fig 3A), as described in details in the materials and methods section and in Fig S3. When the whole cell lysate of this engineered strain was subjected to western blot with anti-mNG antibody, an 82.8 kDa band corresponding to the molecular weight of the fusion protein (31.1 kDa: 3XMyc-mNeonGreen, and 51.7 kDa: Lme-tubulin) was observed. As expected, no band was observed when similar western blot was performed with the lysates from control cells (wild type *L. major* or LmCas9:T7Pol) (Fig 3B).

**Fig 3.**
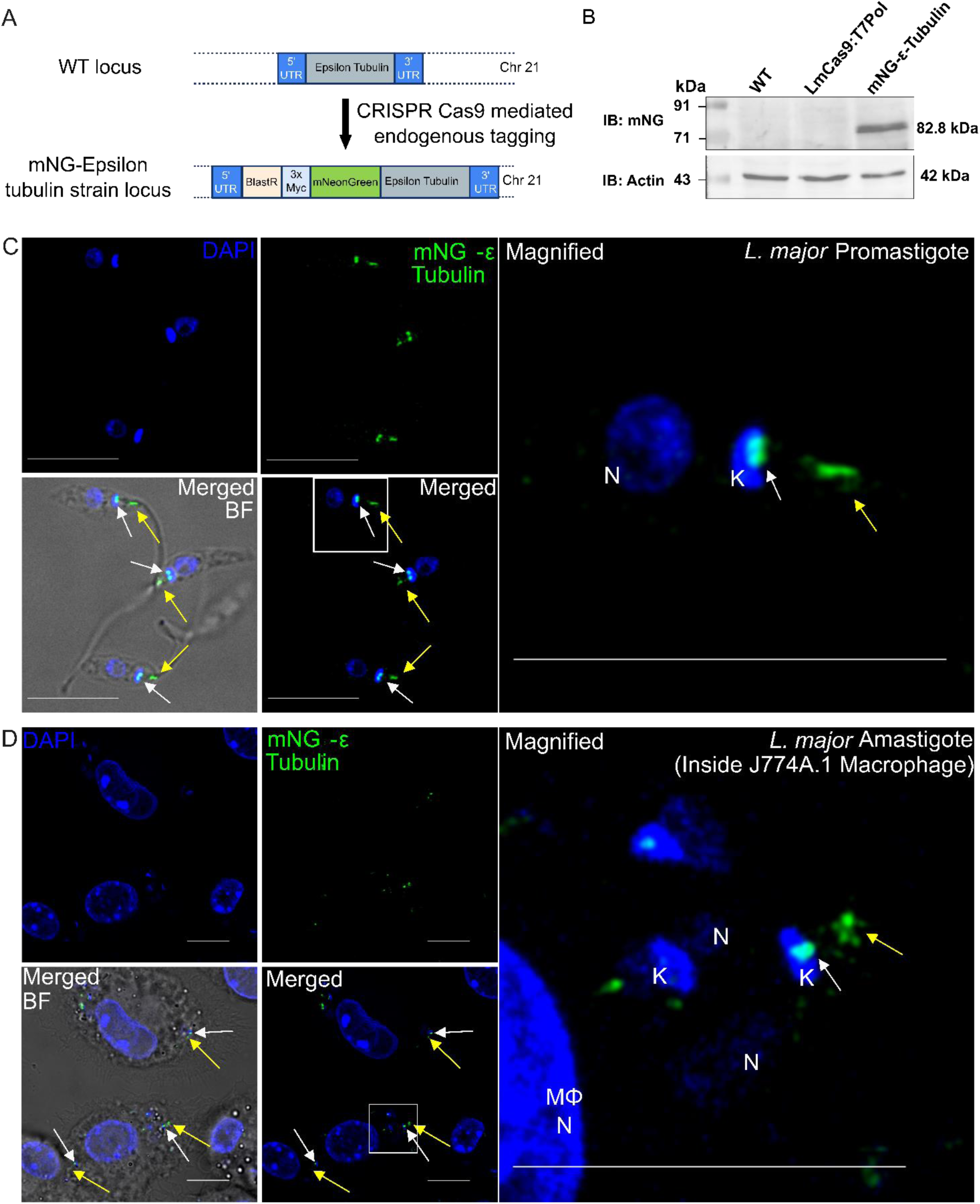
Localization of mNG-tagged epsilon-tubulin in promastigote and amastigote stage *of L. major*. [A] Cartoon diagram showing wild-type (WT) Lme-tubulin genomic locus and engineered mNG-Lme-tubulin genomic locus following CRISPR-Cas9-mediated endogenous tagging. [B] Whole cell lysates from WT, LmCas9:T7Pol and mNG-Lme-tubulin expressing *L. major* strain was subjected to SDS-PAGE and the protein expression was checked by western blotting using antibody against mNG (upper), and actin (lower) as housekeeping control. An 82.8.7 kDa band corresponding to the molecular weight of the fusion protein (31.1 kDa: 3XMyc-mNeonGreen, and 51.7 kDa: Lme-tubulin) was detected. [C] Confocal microscopy of mNG-Lme-tubulin expressing *L. major* promastigotes. Cells were fixed and mounted in DAPI (blue) containing mounting media to stain the nuclei and the kinetoplast. Green signal represents mNG-e-tubulin localization. Merged panel represent the combined image of DAPI along with mNG-e-tubulin. Merged BF represent the fluorescent signal overlay on brightfield image captured simultaneously showing two distinct green signals, one in close association with the kinetoplast DNA and the other at the base of the flagella. The white box represents the area that has been zoomed and represented as magnified panel. The scale bar represents 10μm. The white arrow represents the bright spot of mNG-e-tubulin near kinetoplast (K) and yellow arrow represent bright spot away from kinetoplast. The nucleus is marked as ‘N’ [D] J774A.1 macrophages were infected with mNG-Lme-tubulin expressing *L. major* cells. ∼ 60 hours post infection cells were fixed, and mounted with DAPI-containing mounting media and imaged in a confocal microscope. Parasite nuclei (N) and kinetoplast (K) and macrophage nucleus (MϕN) were stained with DAPI (blue), mNG-e-tubulin is represented by the green signal. Merged image represent the combined image of DAPI along with mNG-e-tubulin. Merged BF represent the fluorescent signal overlay on brightfield image captured simultaneously. The white box represents the area that has been zoomed and represented as magnified panel. The scale bar represents 10 μm. Two bright spots of mNG-e-tubulin can be seen in the amastigotes too, one closely associated with the kinetoplast (white arrow) and the other distal from the kinetoplast (yellow arrow).

### The epsilon-tubulin is localized at two distinct spots in the basal body of *L. major*

After successful validation of the (mNG)-tagged-Lme-tubulin expression *L. major* strain, we aimed to determine intracellular localization of Lme-tubulin in both promastigotes and intracellular amastigotes. Confocal microscopy of the (mNG)-tagged-Lme-tubulin expressing *L. major* promastigotes revealed two bright fluorescent spots in the cell body, one in close association with the kinetoplast DNA and the other at the base of the flagella (Fig 3C). Such a distinct pattern of fluorescent signal indicates basal body localization of Lme-tubulin. The localization pattern of Lme-tubulin is reminiscent of centrin, another basal body-localized protein of the parasite (Selvapandiyan et al., 2001). To check the localization of Lme-tubulin in the intracellular amastigote stage, confocal images were acquired directly from the J774A.1 macrophages infected with the (mNG)-tagged-Lme-tubulin expressing *L. major* strain. Such direct image acquisition method ensures authenticity of the amastigote-character of the parasite, which could otherwise be altered if amastigotes were isolated from the infected macrophages prior to imaging. Like in promastigotes, two distinct fluorescent spots in close proximity of the kinetoplast and the flagellar base were observed in the amastigotes as well (Fig 3D).

To validate basal body localization Lme-tubulin, we treated the (mNG)-tagged-Lme-tubulin expressing *L. major* promastigotes with nocodazole, a known microtubule depolymerizer that does not disrupt the basal body structure. Previous studies in U2OS cells have demonstrated that nocodazole treatment does not affect the localization of human epsilon and delta tubulins, as they are associated with the centrosome and are not part of the microtubular network (Chang and Stearns, 2000; Debrabander et al., 1977). We observed that the localization of Lme-tubulin indeed remained unchanged in >90% of the cells upon treatment with 1 µM nocodazole (Fig 4A, B). To rule out the possibility that nocodazole does not work on *Leishmania* cells, we decided to perform a control experiment to verify if nocodazole treatment could disrupt the localization of α-tubulin, which is an integral part of the microtubular network (McKenna et al., 2023). For this, we generated a mNG-α-tubulin *L. major* strain following similar CRISPR-Cas9-based gene editing method. The authenticity of the strain was confirmed by western blot with anti-mNG antibody, showing a 80.8 kDa band corresponding to the fusion protein (Fig S4). The untreated mNG-α-tubulin in *L. major* promastigotes exhibited characteristic microtubule localization pattern, predominantly along the periphery of the cell, as previously reported earlier in *L. mexicana* (Tran et al., 2015). When these cells were treated with nocodazole, it was observed that unlike the Lme-tubulin, which was largely insensitive to the drug, the α-tubulin localization pattern drastically changed due to microtubule disassembly (Fig 4C, D). Taken together, these data provided unambiguous evidence that Lme-tubulin is indeed localized at the basal body and not part of the broader microtubular network. In this context it is worth mentioning that epsilon tubulin isoforms from *Tetrahymena*, *Chlamydomonas*, *Paramecium* and human have been also shown to localize in the basal body (Chang and Stearns, 2000; Dupuis-Williams et al., 2002; Dutcher et al., 2002; Ross et al., 2013). This suggests that Lme-tubulin may share functional similarity with its counterparts in these organisms.

**Fig 4.**
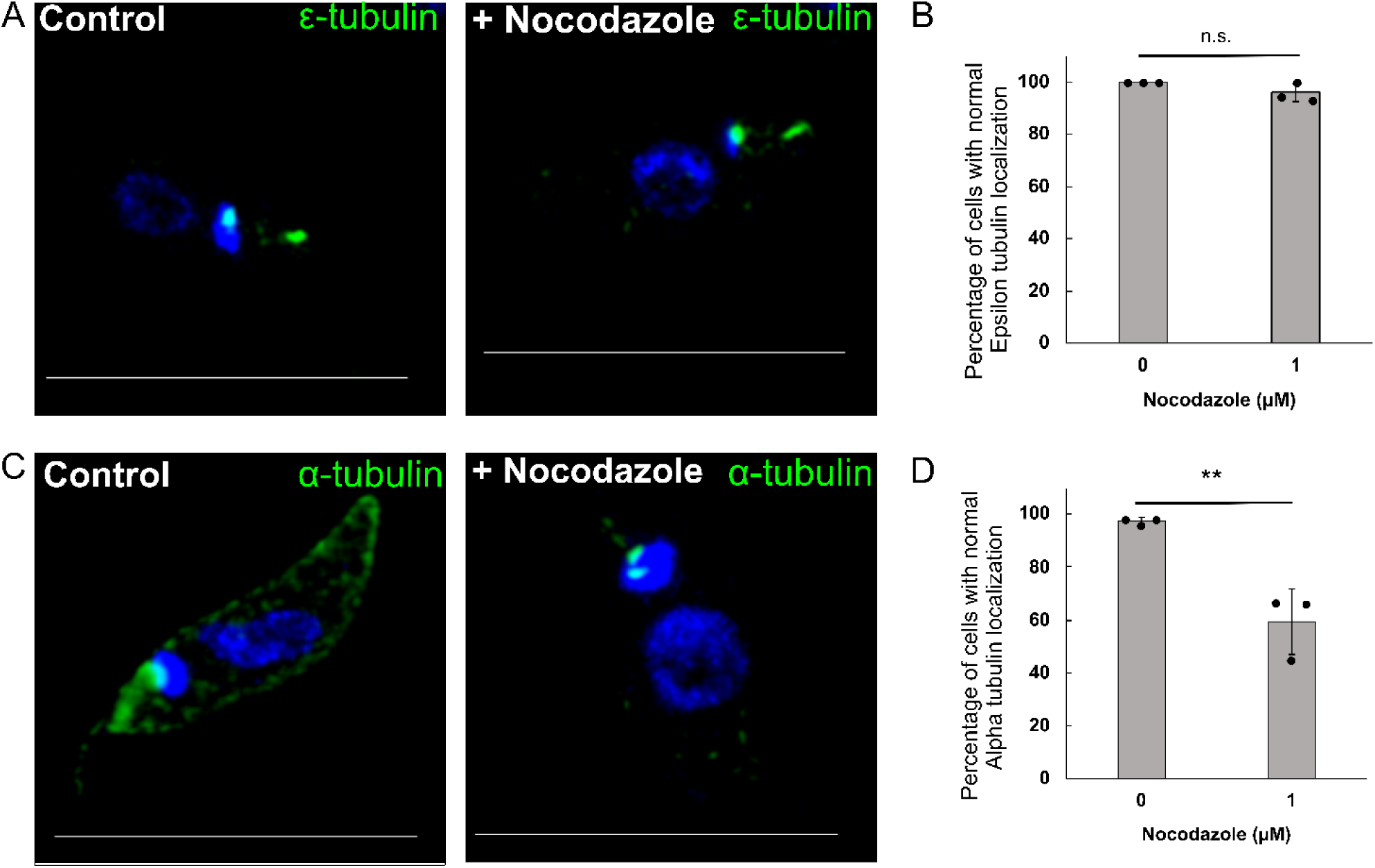
Nocodazole sensitivity assay for tubulins. [A] mNG-ε-tubulin distribution in the untreated (left panel) and 1 μM nocodazole treated (right panel) *L. major* promastigotes expressing mNG-ε-tubulin. Nucleus and kinetoplast are stained with DAPI (blue). Green signal represents mNG-ε-tubulin localization. [B] Bar graph showing percentage of cells where ε-tubulin distribution remained unaltered after mock (only DMSO) or 1 μM nocodazole treatment. [C] mNG-α-tubulin distribution in the untreated (left panel) and 1 μM nocodazole treated (right panel) *L. major* promastigotes expressing mNG-α-tubulin. Nucleus and kinetoplast are stained with DAPI (blue). Green signal represents mNG-α-tubulin localization [D] Bar graph showing percentage of cells where alpha-tubulin distribution remained unaltered after mock (only DMSO) or 1 μM Nocodazole treatment. At least 120 cells were quantified from three independent experiments and presented as ±SD. n.s., non-significant; **p ≤ 0.01.

In conclusion, we identified for the first time a constitutively expressed, basal body-localized e-tubulin not only in the genus *Leishmania* but in the entire *Trypanosomatidae* family. However, further studies, including genetic knockout of Lme-tubulin, are necessary to definitively establish the role of this unconventional tubulin in basal body stability and maintenance. It will be also important to investigate Lme-tubulin’s interaction of with other tubulin isoforms to gain a comprehensive understanding of the microtubule dynamics, especially during promastigote-to-amastigote transition. In particular, investigating the status of ZED module partners of Lme-tubulin, such as delta and zeta-tubulin, would be very interesting to provide deeper insights into the complex interactions that govern the basal body dynamics in *Leishmania* and other trypanosomatid parasites. The CRISPR-Cas9 mediated endogenous protein tagging method employed here to label Lme-tubulin can prove to be very useful for detailed study of the basal body architecture and in identifying different proteins associated with it. Our work is likely to stimulate further research towards these directions.

## Material and Methods

All reagents were purchased from Sigma-Aldrich unless otherwise mentioned. Primers used for PCR were obtained from Integrated DNA Technologies and their sequence information are provided in Table S2. Details of antibodies used in this study are mentioned in Table S3.

### *Leishmania* and mammalian cell culture

*L. major* strain (MHOM/SU/73/5-ASKH) and J774A.1 murine macrophages were procured from ATCC. *L. major* promastigotes were cultured as described before (Pal et al., 2015). Briefly, *L. major* cells were grown at 26 ^0^C in M199 medium (Gibco) supplemented with 15% heat-inactivated fetal bovine serum (Gibco), 23.5 mM HEPES, 0.2 mM adenine, 150 µg/ml folic acid, 10 µg/ml hemin, 120 U/ml penicillin, 120 µg/ml streptomycin, and 60 µg/ml gentamicin. J774A.1 murine macrophages were cultured in Dulbecco’s modified Eagle’s medium (Gibco) pH 7.4, supplemented with 10% heat-inactivated fetal bovine serum, 2 mM L-glutamine, 100 μg/ml streptomycin and 100 U/ml penicillin at 37 °C in a humidified atmosphere containing 5% CO_2_. Whenever necessary, a hemocytometer was used to count cells fixed with 0.9% formal saline or stained with Trypan blue.

### Sequence alignment and bioinformatics

Sequence alignment was performed using the Clustal Omega (https://www.ebi.ac.uk/jdispatcher/msa/clustalo). NCBI Blast (https://blast.ncbi.nlm.nih.gov/Blast.cgi) was used to determine the percent identity and similarity. The result was copied and modified for presentation using MS Word processor and Inkscape without changing any base value. Mutated residues were manually colored as per the literature (Ross et al., 2013) and the tubulin mutation database (https://tubulinmutations.bio.uci.edu/.html). Domains were predicted based on InterPro analysis (https://www.ebi.ac.uk/interpro/). The following protein sequences were obtained from UniProt, epsilon-tubulins from *L. major* (Q4QC95), *T. thermophila* (Q22YZ9), *C. reinhardtii* (A0A2K3DX56), *Homo sapiens* (Q9UJT0), *Paramecium tetraurelia* (Q8WPW7), *L. donovani (A0A3Q8IAD6), L. mexicana* (E9AV85)*, L. braziliensis* (A4HBW1)*, L. infantum* (E9AGW9)*, T. cruzi* (A0A7J6XYD9)*, T. brucei* (Q9NI44) *L. major* α-tubulin (Q4QGC5), *L. major* β-tubulin (Q4Q4C4), *L. major* γ-tubulin (Q4Q9Z8). The predicted structure for *L. major* α-, β- and γ-tubulin was obtained from the AlphaFold server (Jumper et al., 2021; Varadi et al., 2022). Structural alignment was done using the ChimeraX software module (Pettersen et al., 2004).

### Genomic DNA PCR

The genomic DNA of *L. major* promastigotes was isolated using the phenol-chloroform method. Briefly, 1 x10^7^ *L. major* cells were harvested and resuspended in 0.1 ml lysis buffer (0.1 M Tris-Cl, 0.1 M NaCl, 1% SDS, 100 mM EDTA, 100 μg/ml Proteinase K) and incubated at 55 ^0^C for 120 minutes. 25 μl phenol and 25 μl chloroform were added to the lysate and centrifuged at 13000g for 15 minutes. The upper aqueous phase was taken in a fresh tube, and an equal volume of chloroform was added and centrifuged at 13000g for 15 minutes. The supernatant was collected, and 1/10 volume 3 M sodium acetate and 2 volume ethanol were added and centrifuged at 13000g for 10 minutes. The pellet is washed with 70% ethanol twice and eluted in nuclease-free water. DNA Concentration was determined spectrophotometrically using NanoDrop spectrophotometer. Genomic DNA PCR amplification was performed using Leo Mastermix (Dxbidt, India) and specific primers following manufacturer’s protocol.

### Checking of Lme-tubulin expression in *L. major* promastigote

Total RNA from *L. major* promastigotes was isolated using Trizol reagent (Invitrogen) followed by DNase I treatment (Invitrogen). cDNA was prepared from DNase I-treated RNA using Verso cDNA kit (Thermo Scientific) using the manufacturer’s protocol. -RT controls were prepared from DNase I treated RNA where all Verso cDNA kit components were added except the reverse transcriptase enzyme. PCR was performed using the cDNAs from +RT and -RT samples.

### Checking of Lme-tubulin expression in *L. major* amastigote

*L. major* infection of J774A.1 macrophages was performed as described by us before (Pal et al., 2017). Briefly, J77A.1 macrophages were activated in the presence of 100 ng/ml LPS for 6 hours, and to it stationary phase *L. major* cells were added at MOI 1:30. After 12 hours, media was discarded to remove uninternalized parasites, and washed with 1X PBS, and fresh DMEM media was added, and incubated for ∼60 hours. Total RNA was extracted from infected and uninfected macrophages and RT-PCR was performed following the protocol described above.

LmCas9:T7Pol *L. major* strain generation

LmCas9:T7Pol expressing *L. major* cells were generated following protocols and plasmid developed by Beneke et al. (Beneke et al., 2017) with slight modification. pTB007 cells were electroporated into wild-type *L. major* cells. 30 μg of the plasmid was resuspended in an electroporation buffer (21 mM HEPES, 0.7 mM NaH_2_PO_4_, 137 mM NaCl, and 6 mM glucose; pH 7.4) and electroporated using BTX Gemini electroporator (480 V, 550 µF, 25 ohms) to 4 x10^7^ *L. major* promastigotes. The cells were selected in the presence of 50 μg/ml Hygromycin. Total genomic DNA was isolated from selected cells and subjected to PCR to verify the generated strain. The PCR was conducted with P3/P4, P5/P6 and P7/P8 for amplification of SpCas9, T7pol and rRNA45, respectively.

### Generation of N-terminal mNG-tagged-epsilon/alpha-tubulin expressing *L. major* strains

The N-terminal mNG-tagged-epsilon/alpha-tubulin expressing *L. major* strains were generated following the protocol reported by Beneke et al. (Beneke et al., 2017) and as schematically described in Fig S3. The primers used for the strain generation were designed using the LeishGEdit primer design module (http://www.leishgedit.net/Home.html) and the primer sequences are provided in Table S2. For preparation of sgDNA (template for sgRNA), overlap extension PCR was performed using primer P11 (for ε-tubulin) or P14 (for α-tubulin) and primer G00 (containing sgRNA scaffold sequence). Donor DNA containing blasticidin resistant gene flanked by 30 nucleotide upstream and downstream sequence of the Cas9-cleavage site was PCR amplified using pPLOTv1 blast-mNG-blast plasmid as template and primers P9/P10 (for ε-tubulin) and P12/P13 (for α-tubulin). All the PCR amplifications were done using PFU polymerase following the protocol mentioned by Beneke et al., 2017 (Beneke et al., 2017). The donor DNAs were gel eluted using commercial kit (BioBharati). The eluted donor DNA and corresponding sgDNA were simultaneously electroporated into LmCas9:T7Pol *L. major* cells and selected in 20 μg/ml blasticidin S containing media for ∼7 days.

### Western blotting

Whole cell lysate was prepared in 1X resuspension buffer (1X PBS+1X protease inhibitor cocktail) with sonication. Protein amount was quantified using Lowry method and 40 µg of whole cell lysate were loaded in 12% polyacrylamide gel and subjected to SDS-PAGE. The protein was transferred to PVDF membrane and blocked with blocking solution (5% skim milk solution in TBST (10 mM Tris, 150 mM NaCl, 0.025% Tween 20)). The membrane is washed for 5 minutes for two times and incubated overnight with shaking at cold room with rabbit anti-mNG primary antibody and rabbit anti-LdActin diluted in TBST (Table S3). The membrane was washed with 1X TBST for 5 minutes 5 times and incubated with HRP tagged goat anti-rabbit secondary antibody for 1.5 hours in room temperature. The membrane was washed with 1X TBST for 5 minutes for 5 times. Using chemiluminescent substrate (Biorad), blot was developed and signal documented in ChemiDoc imaging system (Biorad).

### Imaging studies

mNG-ε-tubulin expressing *L. major* promastigotes were spread on poly-L-lysine-coated coverslips and incubated for 30 minutes. Fixation solution (4% paraformaldehyde in 1X PBS) was added to it and incubated in dark for 10 mins. The coverslips were washed twice with 1X PBS and mounted in antifade mounting media with DAPI (Vectashield). The images were acquired in Leica SP8 confocal microscope using a 63X oil immersion lens. Images were processed using LasX (Leica). Central 1-3 stacks were merged and projected for representation. mNG-ε-tubulin expressing *L. major* amastigote inside macrophages were imaged by the same protocol with slight modification. J774A.1 macrophages were infected with mNG-tagged-ε-tubulin *L. major* cells as described above. At ∼60 hours post infection, macrophages were fixed, washed twice with 1X PBS and mounted in antifade mounting media with DAPI (Vectashield). The images were acquired in Leica SP8 confocal microscope using a 63X oil immersion lens. Images were processed using LasX (Leica). Central 1-3 stacks were merged and projected for representation.

### Nocodazole disruption assay

mNG-tagged-epsilon/alpha-tubulin expressing *L. major* promastigotes (1x10^7^ cells) were incubated with 1 μM Nocodazole or DMSO (as control) in 200 μl M199 media for 1 hour. Cells were spread on a poly-L-lysine coated coverslip and fixed with 4% paraformaldehyde for 10 minutes. Coverslips were washed twice with 1X PBS, mounted in antifade mounting media with DAPI (Vectashield) and imaged immediately in Leica SP8 confocal microscope. Individual stacks were analyzed and represented. The percentage of cells with unaltered localization of the respective tubulins (ε or α) were calculated for both untreated and nocodazole treated cells. In the case of α-tubulin, the cytoskeleton is considered intact if the peripheral tubulin signal is clearly visible. In the case of ε-tubulin strain, if two distinct puncta are visible, it is considered as unaltered.

## Acknowledgment

The authors thank Dr. Sankar Maiti and Dr. Subhra Majumder for sharing reagents and for providing valuable suggestions. Dr. Subrata Adak and Dr. Amogh Sahasrabuddhe are acknowledged for providing the plasmids for CRISPR-Cas9 manipulation and LdActin antibody, respectively. Dr. Subhankar Dolai is thanked for his critical comments on the manuscript.

## Data and resource availability

All other data that supports the results of this study are available from the corresponding author upon reasonable request.

## Supplementary information

**Fig. S1.**
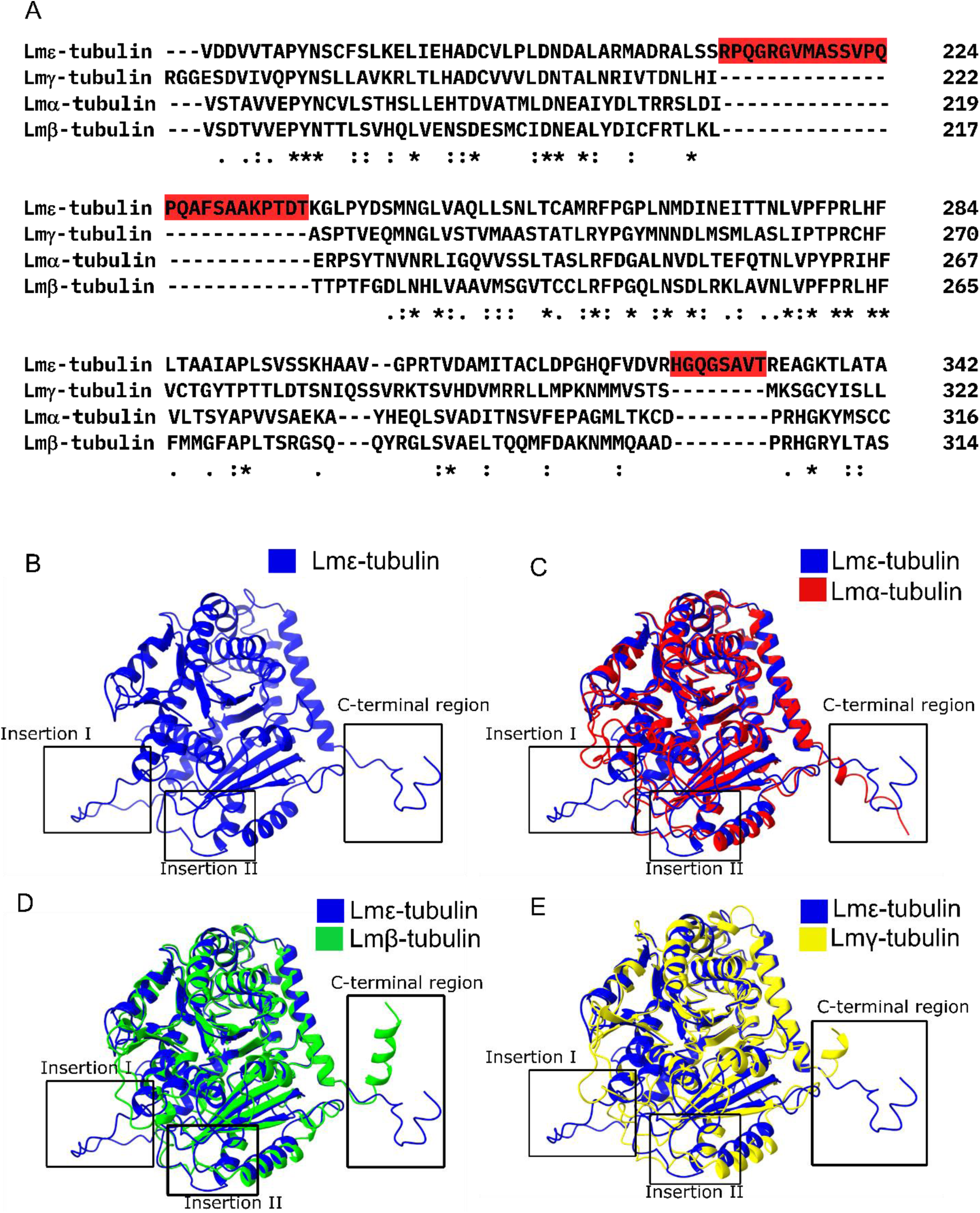
Sequence and structural alignment of Lmε−tubulin with other conventional tubulins of *L. major*. [A] Lmε-tubulin (168-342) protein sequence was aligned with Lmα-tubulin (177-316), Lmβ-tubulin (175-314) and Lmγ-tubulin (177-322). Although the complete sequence of all the tubulins were aligned, only a portion of the sequence (the exact region mentioned in parenthesis) are represented in the figure. The regions shaded red indicate insertions in Lmε-tubulin that are absent in α, β or γ-tubulins, Insertion I-R211-T236, Insertion II-H325-T332. [B] AlphaFold-predicted structure of Lmε-tubulin, Insertion I, Insertion II and the C-terminal region are marked with boxes. Structural alignments of AlphaFold structure of Lmε-tubulin with that of AlphaFold structures of [C] Lmα-tubulin, [D] Lmβ-tubulin, [E] Lmγ-tubulin. All the structures aligned well with Lmε-tubulin except for three regions, Insertion I, Insertion II and the C-terminal tail.

**Fig. S2.**
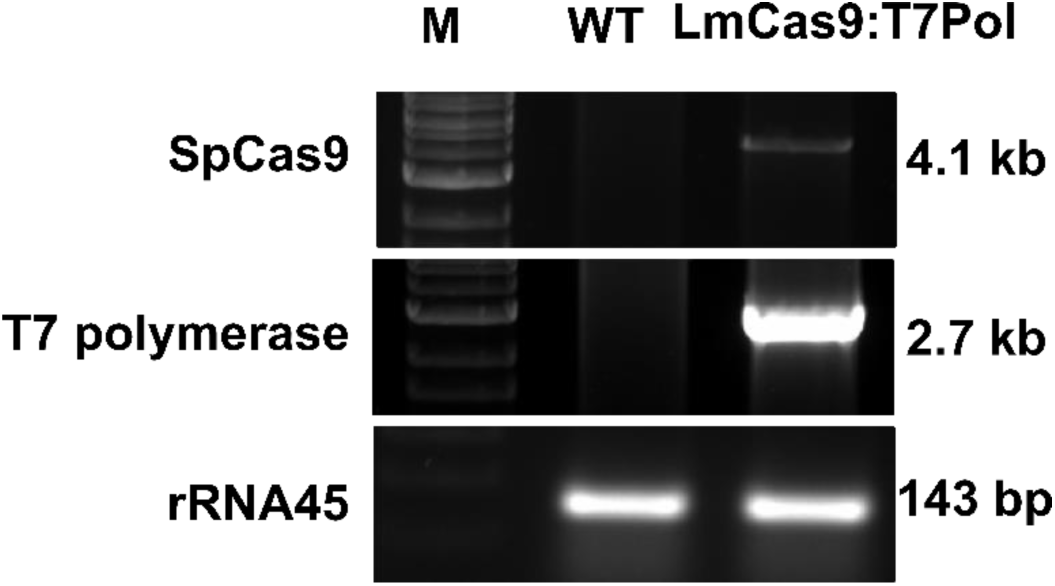
Verification of LmCas9:T7Pol *L. major* strain. Presence of SpCas9 and T7 polymerase gene in the genome of LmCas9:T7Pol *L. major* cells was checked by PCR. PCR was performed with genomic DNA from wild-type (as negative control) or LmCas9:T7Pol *L. major* cells as templates with specific primers for SpCas9 and T7 Polymerase. 4.1 kb (for SpCas9) and 2.7 kb (for T7 Pol) PCR products in the LmCas9:T7Pol *L. major* cells but not in the WT confirmed proper generation of the strain. PCR for the housekeeping gene rRNA45 was used as loading control.

**Fig. S3.**
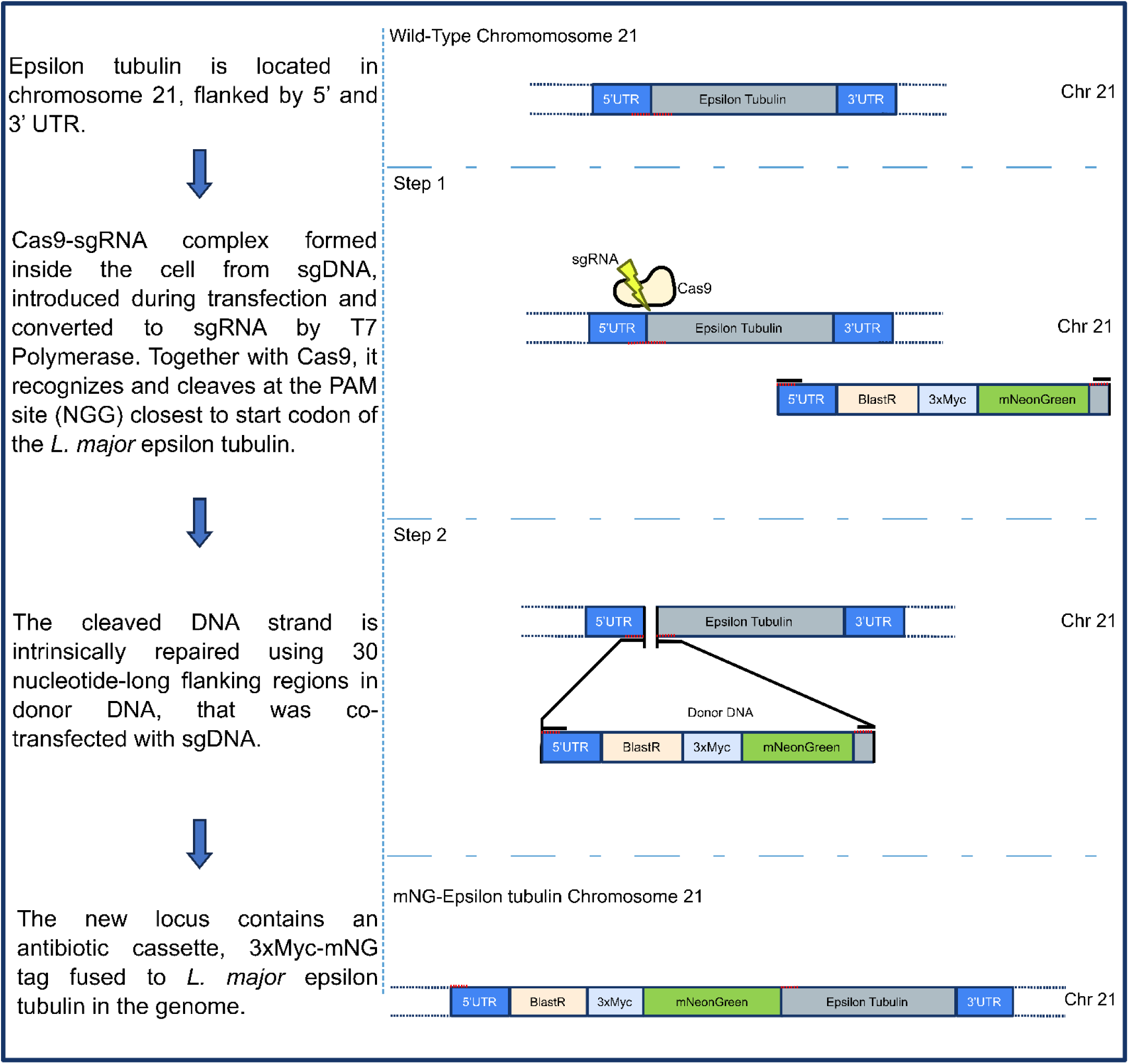
Scheme for generation of mNG-Lmε-tubulin expressing *L. major* strain. Steps for generation of the N-terminal mNG-tagged-Lmε-tubulin expressing *L. major* strain employing CRISPR-Cas9 endogenous tagging method are described schematically.

**Fig S4.**
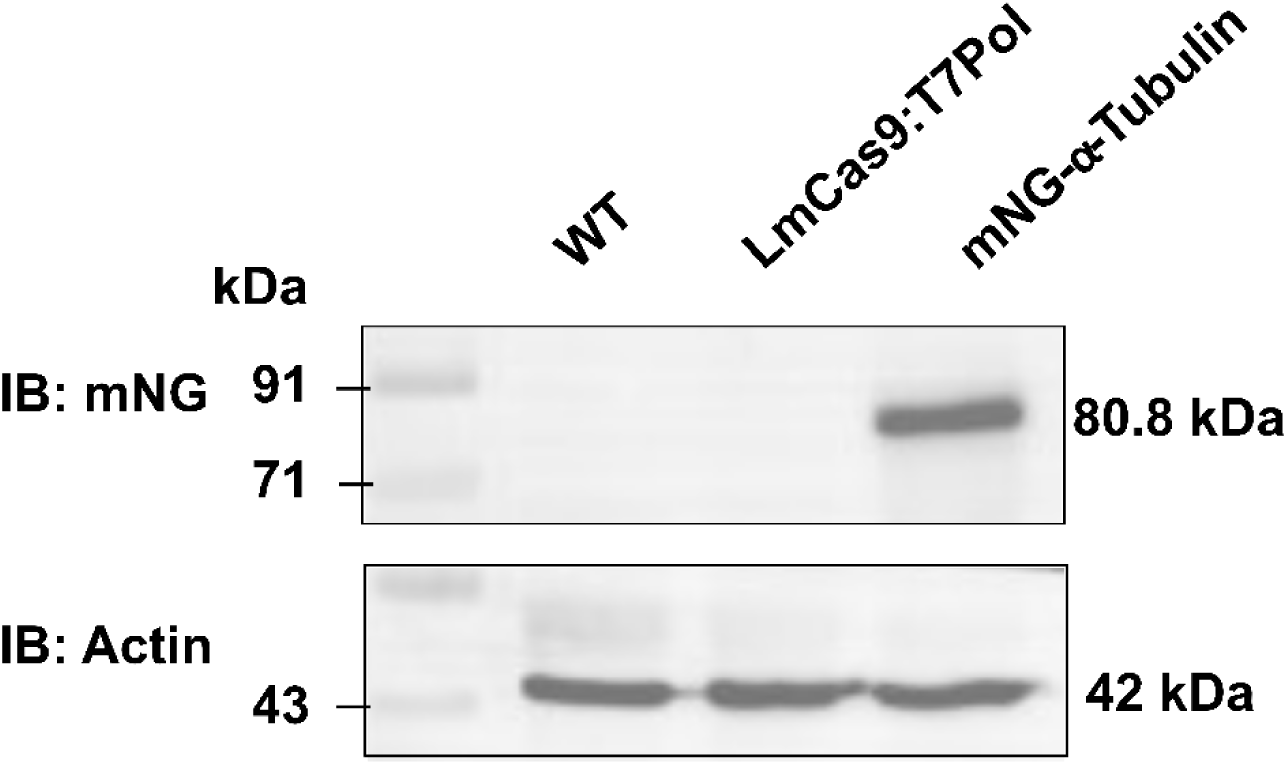
Confirmation of the mNG-Lmα-tubulin expressing *L. major* strain by western blot. Whole cell lysate from wild-type (WT), LmCas9:T7Pol and mNG-Lmα-tubulin expressing *L. major* strain was subjected to SDS-PAGE and the protein expression was checked by western blotting using antibody against mNG (upper), and actin (lower) as housekeeping control. A 80.8 kDa band corresponding to the molecular weight of the fusion protein (31.1 kDa: 3XMyc-mNeonGreen, and 49.7 kDa: Lmα-tubulin) was detected.

**Table S1.** Percentage identity and similarity of Lme-tubulin with other organisms.

| **Genus** | **Species** | **Percentage identity with *L. major* (%)\*** | **Percentage similarity with *L. major* (%)\*** |
| --- | --- | --- | --- |
| *Tetrahymena* | *T. thermophila* | 40.97 | 59.43 |
| *Chlamydomonas* | *C. reinhardtii* | 43.83 | 60.08 |
| *Homo* | *H. sapiens* | 41.37 | 59.04 |
| *Paramecium* | *P. tetraurelia* | 41.65 | 61.31 |
| *Leishmania* | *L. donovani* | 98.72 | 99.15 |
|  | *L. infantum* | 98.72 | 99.15 |
|  | *L. mexicana* | 97.23 | 98.51 |
|  | *L. braziliensis* | 91.06 | 95.96 |
| *Trypanosoma* | *T. brucei* | 68.24 | 80.12 |
|  | *T. cruzi* | 66.80 | 82.35 |
| **\*Percentage identity and similarity calculated using NCBI blast** |  |  |  |

**Table S2.**
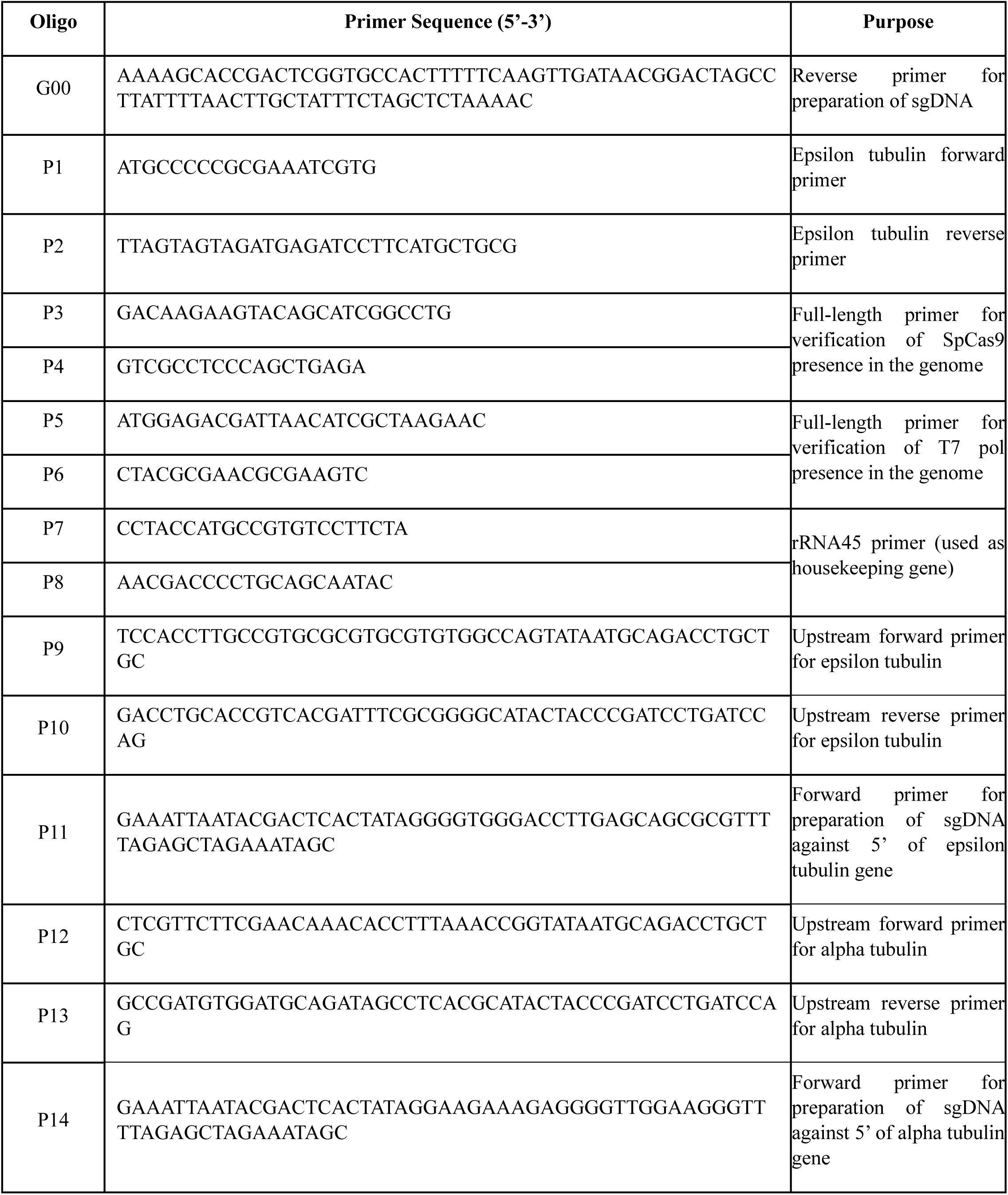
Oligonucleotides used in this paper.

**Table S3.** Antibodies used in this study.

| **Antibody** | **Host** | **Source** | **Catalogue Number** | **Dilution** |
| --- | --- | --- | --- | --- |
| mNeonGreen Tag Antibody | Rabbit | Cell Signaling Technology | 53061S | WB- 1:1000 |
| Anti-Ld Actin | Rabbit | Gifted by Dr. Amogh Sahasrabuddhe (CSIR-CDRI) | - | WB- 1:20000 |
| Goat Anti-Rabbit IgG-Peroxidase | Goat | Sigma Aldrich | A9169 | WB- 1:4000 |

